# Development and characterization of a multimeric recombinant protein based on the spike protein receptor binding domain of SARS-CoV-2 that can neutralize virus infection

**DOI:** 10.1101/2023.02.15.528632

**Authors:** Veronica Aparecida de Lima, Rodrigo da Silva Ferreira, Maria Luiza Vilela Oliva, Robert Andreata-Santos, Luiz Mario Ramos Janini, Juliana Terzi Maricato, Milena Apetito Akamatsu, Paulo Lee Ho, Sergio Schenkman

**Affiliations:** Department of Microbiology, Immunology and Parasitology, Escola Paulista de Medicina, Universidade Federal de São Paulo, São Paulo, Brazil; Department of Biochemistry, Escola Paulista de Medicina, Universidade Federal de São Paulo, São Paulo, Brazil; Núcleo de Produção de Vacinas Bacterianas, Centro BioIndustrial, Instituto Butantan, São Paulo SP, Brazil

**Keywords:** SARS-CoV-2, COVID-19, Neutralizing antibody, Protein S, Receptor-binding domain

## Abstract

**Background:** The SARS-CoV-2 virus, responsible for the COVID-19 pandemic, has four structural proteins and sixteen non-structural proteins. The S-protein is one of the structural proteins exposed on the surface of the virus and is the main target for producing neutralizing antibodies and vaccines. The S-protein forms a trimer that can bind the angiotensin-converting enzyme 2 (ACE2) through its receptor binding domain (RBD) for cell entry.

**Methods:** We stably expressed in a constitutive manner in HEK293 cells a new recombinant protein containing a signal sequence of immunoglobulin to produce an extended C-terminal portion of the RBD followed by a region responsible for the trimerization inducer of the bacteriophage T4, and a sequence of 6 histidines. The protein was produced and released in the culture supernatant of cells and was purified by Ni-agarose column and exclusion chromatography. It was then characterized by SDS-polyacrylamide gel and used as antigen to generate protective antibodies to inhibit ACE2 receptor interaction and virus entry into Vero cells.

**Results:** The purified protein displayed a molecular mass of 135 kDa and with a secondary structure like the monomeric RBD. Electrophoresis analysis in SDS-polyacrylamide gel with and without reducing agents, and in the presence of crosslinkers indicated that it forms a multimeric structure composed of trimers and hexamers. The purified protein was able to bind the ACE2 receptor and generated high antibody titers in mice (1:10000), capable of inhibiting the binding of biotin labeled ACE2 to the virus S1 subunit, and to neutralize the entry of the SARS-CoV-2 Wuhan strain into cells.

**Conclusion:** Our results characterize a new multimeric protein based on S1 subunit to combat COVID-19, as a possible immunogen or antigen for diagnosis.

## Introduction

In December 2019, patients in Wuhan, the capital of China’s Hubei Province, experienced symptoms of severe influenza, mainly affecting the respiratory tract. They were initially reported to the Wuhan Municipal Health Department as unexplained cases of pneumonia. At the same time, the Wuhan Center for Disease Control and Prevention discovered that these patients were associated with a seafood market [1]. Sequencing of samples isolated from the patients confirmed that the pathogen was a new type of coronavirus. As cases increased worldwide, human-to-human transmission was identified as the main cause of transmission. Epidemiological studies reported the occurrence of infection in people who did not have access to the seafood market in Wuhan [2]. On February 11, 2020, the World Health Organization designated the disease caused by the SARS-CoV-2 virus as Coronavirus Disease-2019 (COVID -19), and on March 20, 2020, it designated COVID-19 as a pandemic. By December 2022, there were more than 657 million confirmed cases and more than 6.6 million deaths worldwide (https://www.who.int/emergencies/diseases/novel-coronavirus-2019/situation-reports).

Coronaviruses belong to the *Coronaviridae* family of the subfamily *Coronavirinae*, which includes four genera: Alphacoronavirus, Betacoronavirus, Gammacoronavirus, and Deltacoronavirus [3]. There are seven species of human coronaviruses (HCoV), four of which circulate and usually cause mild infections; HCoV-29E, HCoV-OC43, HCoV-NL63, and HCoV-HKU1 [4], and three that cause more severe respiratory symptoms; SARS-CoV, MERS-CoV and SARS-CoV-2. SARS-CoV-2 has approximately 80% similarity to SARS-CoV and 51.8% similarity to MERS-CoV [5], but the closest genetic identity of SARS-CoV-2 is SARS-RaTG13 bat CoV with approximately 96.2% compatibility (Hu et al., 2021). The evolutionary ability of coronaviruses has been recognized, and this has an implication on the development of new diagnostic processes, as large number of new coronavirus genomes are appearing continuously [6].

SARS-CoV-2 is a betacoronavirus whose genome is a positive single-stranded RNA that varies in size from 27 to 32 kbases. It contains four structural proteins and sixteen non-structural proteins [3]. These structural proteins are spike proteins (S), a transmembrane homotrimeric glycoprotein formed by two subunits, S1and S2. The S1 subunits are responsible for viral entry into the host cell through its receptor binding domain (RBD), which binds to angiotensin converting enzyme 2 (ACE2) in the host cell. The S2 subunit is responsible for viral and host cell membrane fusion [7].

Protein S is the most important target for vaccine development as binding to protein S generates neutralizing effects, which prevent the interaction with the cell receptors [8]. For this reason, there are several studies using the whole S protein with minor modifications as a vaccine candidate [9] and several monoclonal antibodies have been obtained [10]. However, the generation of neutralizing antibodies requires maintenance of the protein in state prior to fusion processing occurring during virus cell fusion [11-13]. Alternatively, protective antibodies can be generated by immunization with RBD, which is essential for SARS-CoV-2 entry into the host cell [14]. The RBD contains an extended insert, the receptor binding motif with amino acid residues that bind to the N-terminus of ACE2 [15]. Multimeric RBDs have been shown to be more efficient at generating neutralizing antibodies because they can increase B-cell activation [16], enhance neutralizing antibody responses, and prevent the formation of low-affinity antibodies [17]. Therefore, several works have shown that trimeric RBD induces higher titers of neutralizing antibodies compared to monomeric RBD [18]. For instance, immunization of *rhesus macaques* with various forms of protein S showed protective humoral and cellular responses, in addition to neutralizing antibody titers (J. Yu et al., 2020).

Today, there are more than 400 preparations in clinical development and pre-clinical development (https://www.who.int/publications/m/item/draft-landscape-of-covid-19-candidate-vaccines). The main available vaccines, Moderna (mRNA-1273) and Pfizer/BioNtech (BNT162b2) use mRNA, which encodes glycoprotein S and produces high titers of neutralizing antibodies with 94 to 95% efficacy [19-22]. The vaccine ChAdOx1 nCoV-19 (AZD1222) consists of a chimpanzee adenoviral vector with a replication defect, which contains the glycoprotein gene S as an antigen. This vaccine showed an efficacy of 70% [23]. The Ad26.COV2.S vaccine produced by Johnson & Johnson consists of a recombinant adenovirus serotype 26 (rAd26) that also encodes protein S [24]. It has been proven to be effective after a single dose with high humoral response [25]. Another vaccine that uses the adenoviral vector rAd26 and rAd5 is Sputnik V, which has been shown to induce strong humoral and cellular responses with two doses [26]. The inactivated virus is also used in the production of the CoronaVac vaccine that uses SARS-CoV-2 in its inactive form [27] and was able to produce a strong immune response [28].

Although numerous studies have generated proteins with antigenic properties of the SARS-CoV-2 RBD, our group has developed an antigen that in addition to the RBD domain, included a C-terminal portion of the S1 protein linked to the trimerization region of bacteriophage T4 in the C-terminus of the protein to generate a robust and stable antigen expression for diagnostics, immunization, and neutralization assays. In contrast to other studies, we generate cells stably, expressing the protein to allow a simple production and purification of a trimeric form of the RBD antigen.

## Material and Methods

### Animals

Female BALB/c mice aged 6-8 weeks, obtained from the Centre for the Development of Experimental Models for Medicine and Biology, CEDEME – Universidade Federal de São Paulo (UNIFESP), were used. Animals were maintained under standard lighting conditions (12 hours of light and 12 hours of darkness) at controlled temperature (25 ± 2 ºC) and with food and water ad libitum in the vivarium of the Department of Microbiology at the Escola Paulista de Medicina of UNIFESP. For euthanasia, animals received a lethal dose of xylazine and ketamine intraperitoneally in accordance with ethics committee regulations. All procedures in this work were approved by the Ethics Committee on the Use of Animals of UNIFESP (Case No. 8507290721).

### Plasmid and recombinant production

The pcDNA 3.1 (+) plasmid containing the RBD sequence of the protein (encompassing the region between amino acids 331 to 589 of the SARS-CoV-2 S protein (MT350282) was synthesized by GeneScript after codon optimization for expression in mammalian cells. The insert DNA sequence is shown in the supplementary **Fig**. S1. The plasmid was grown in *E. coli* Mach1 (provided by the Structural Genomic Consortia, SGC Campinas, Brazil) in Luria-Bertani medium containing 100 µg/mL ampicillin (Sigma-Aldrich). Plasmid DNA was prepared using the Midi-Prep kit (Sigma-Aldrich) and preparations were analyzed after digestion with endonucleases by electrophoresis in 0.8% agarose gel in TAE buffer (40 mM Tris-HCl, 0.3 mM acetic acid and 2 mM EDTA).

### Cell culture

Human cells (HEK293) (Human embryonic kidney 293, ATCC CRL -1573), kindly provided by Prof Dr João Bosco Pesquero, UNIFESP, were cultured with RPMI 1640 (Thermo Fisher Scientific), 50 U/mL penicillin, and 50 µg/mL streptomycin, supplemented with 10% FBS (InvitroCell, Brazil), and maintained at 37ºC and 5% CO2. For transfections, 2 × 10^4^ HEK293 cells were added to each well of a 6-well plate, and after 24 hours at 37 °C, cells were transfected with the linearized plasmid DNA-lipofectamine LTX complex (Thermo Fisher Scientific) according to the manufacturer’s recommendations. 0.5 µg of plasmid pcDNA 3.1/RBD was linearized with BamHI enzyme (Thermo Fisher Scientific), precipitated, and washed with 70% ethanol. The DNA was resuspended with 285 µl Opti-MEM (Thermo Fisher Scientific). Then, 15 µl of Lipofectamine LTX (Thermo Fisher Scientific, USA) was added and the solution was incubated for 20 min at room temperature to form the DNA-lipofectamine complex. The cell monolayers were washed once with Opti-MEM, and 0.5 ml of Opti-MEM, containing the plasmid, was added to each well. Cells were incubated at 37°C, and after 24 hours, the medium was replaced, and selection was initiated with 0.5 mg/mL Geneticin-G418 (InvivoGen, USA). Plasmid-free transfected cells were used as a control. To confirm secretion of the recombinant protein RBD, cells were cultured with RPMI and 10% FBS until they reached 90% confluence. After reaching confluence, cells were cultured with medium without FBS (Gibco, USA), and the supernatant was collected for 4 days for analysis by Western blotting.

### Western blotting and immunofluorescence

For Western blotting assays, samples were boiled with sample buffer (50 mM Tris-HCl, pH6.8, 0.02% bromophenol blue, 2% SDS, and 20% glycerol, with or without 10% β-mercaptoethanol), applied to a 10% SDS-PAGE gel, and transferred to 0.45-µm nitrocellulose membranes (Bio-rad, USA). The membranes were blocked with 5% BSA or 5% skim milk powder in 0.15 M NaCl, 10 mM Tris-HCl pH 7.4 (TBS), for 1 hour. The membranes were then incubated for 1 hour with horse serum from horses immunized with Newcastle virus-expressing spike antigen at a dilution of 1:10.000 in blocking solution or with serum from mice immunized with a recombinant RBD protein produced in bacteria (kindly provided by Dr. Santuza Teixeira from the Federal University of Minas Gerais). The sera from mice immunized with the recombinant protein produced in this work was also used. After washing the membranes with TBS, which contained 0.05% Tween 20 (TBS-T), binding of equine antibodies was detected by incubation for 1 hour with protein A conjugated to peroxidase (GE) and visualized with chemiluminescence reagent (ECL, Merck). Images were obtained with an Odyssey LI-COR instrument. Mouse antibodies were detected by binding to IRDye800 mouse anti-IgG (LI-COR) and observed with the same system.

For immunofluorescence, transfected cells (2 × 10^4^) were plated onto 13-mm glass coverslips in 24-well plate. After 24 hours they were fixed with 4% p-formaldehyde (PFA) in PBS for 20 minutes on ice. Fixed cells were washed three times with PBS permeabilized with 0.05% Triton X100 in PBS for 2 minutes, washed again in PBS and maintained for 30 min with 3% BSA in PBS before incubation with the indicated serum. Bound antibodies were detected by adding Alexa488 conjugated to anti-IgG (1:10000, Thermo Fisher Scientific) and then observing them in an Olympus BX61 fluorescence microscope.

### Expression and purification of the recombinant protein

For protein production, aliquots of cells frozen in liquid nitrogen were grown in 75 cm^2^ culture flasks in RPMI 1640 medium containing 10% FBS and 0.5 mg/ml Geneticin G418 until confluence was reached. Cells were trypsinized and seeded in four 150cm^2^ flasks containing the same culture medium. After reaching 80% confluence, the medium was replaced by 25 ml of medium without FBS (OptiPro SFM, 1230919, Thermo Fisher Scientific) and geneticin G418. The medium was replaced after 4 days, and protein expression in the supernatant was monitored by Western blotting. Culture supernatants that contained detectable recombinant protein were collected and stored at 4°C until purification.

Protein purification was performed by loading the culture supernatant at 1 ml/min onto a Ni-Sepharose HisTrap FF™ column (GE, USA) pre-equilibrated with 40 mM imidazole, 20 mM sodium phosphate, and 0.5 M NaCl, pH 7.4. In each batch, 400 ml of supernatant was used. After one passage, the column was washed with equilibration buffer and the protein was eluted with 15 ml of 0.5 M imidazole, 20 mM sodium phosphate, 0.5 M NaCl, pH 7.4 in three fractions of 5 ml. These fractions were concentrated by centrifugation on Centricon Plus-20 5,000 filters (Merck-Millipore), and protein levels were determined by micro-BCA assays (Thermo Fisher Scientific).

The concentrated samples (0.5 mL) were chromatographed on a Superdex 200 column (30 × 1, Cytiva) equilibrated in TBS at a flow rate of 0.4 mL/min. 0.25 mL fractions were collected using an Akta purifier chromatographer (GE). For determination of molecular mass, the column was calibrated with BSA (66 kDa and 132 kDA), carbonic anhydrase (30 kDa), and β-amylase (200 kDa) (Sigma-Aldrich).

### Structural analysis

CD measurements were performed using a Chirascan Plus spectrometer (Applied Photophysics, Leatherhead, UK) at 25°C; optical path of 0.1 cm, in quartz cuvettes. CD data were obtained in the range of 200-250 nm, with 1 nm step, and 1 nm window. Protein concentrations used were 0.345 mg/mL (2.5 µM), and spectra were obtained from the average of 8 scans. Spectra were converted to molar ellipticity [θ] using Equation 1 for secondary structure analysis and estimated using CDPro software [29].

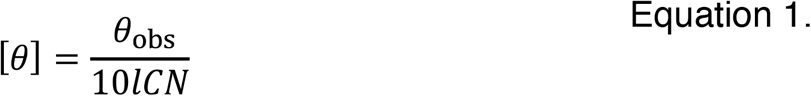

Where *θ*_obs_ is the CD in millidegrees, C is the protein concentration (M), l is the path length of the cuvette (cm), and N is the number of amino acid residues.

Cross-linking experiments were performed by incubation of 2 mM and 5 mM of suberic-bis-acid ester (3-sulfo-N-hydroxyuccinimy (BS3, Sigma-Aldrich) with the purified recombinant protein in 25 mM sodium phosphate pH 7.4 for 1 hour. To stop the reaction, 25 mM of Tris-HCl pH 7.4 were added, and the mixture incubated for another 10 minutes. The samples were analyzed by Western blotting. Structural model was performed using the Chimera software [30].

### Immunization and immunoassays

Mice were divided into two groups (n=5 per group), one experimental group (immunized with RBD-T) and one control group (PBS). Animals were immunized intramuscularly in the hind paw with 100 µg of the purified RBD protein diluted in PBS 1x in the presence of the same volume of aluminium hydroxide (Inject Alum, Thermo Fisher Scientific). A total of 3 doses of 0.05 µL were administered into each paw (Prime, Boost I, and Booster II), two weeks apart. Blood sampling to check antibody titers was performed via the tail vein (10 µL of blood), 15 days after each dose. Fifteen days after the last dose, animals were anesthetized, and blood was collected by cardiac puncture.

For ELISA assays, a white high-binding polystyrene plate (Sarstedt) was used, sensitized by the addition of 200 ng of the purified RBD protein diluted in 50 µL of 0.1 M sodium bicarbonate pH 8.5 per well, and incubated overnight at 4°C. The wells were washed twice with PBS-Tween 0.05%, (PBS-T) and were incubated with 5% skim milk powder in PBS for 1 hour and then with the sera diluted in the same solution for 1 hour at 37°C. The wells were washed again in PBS-T and incubated with 100 µL of 1/10000 diluted anti-mouse peroxidase (Thermo Fisher Scientific). After one hour, the wells were washed, and binding was detected by the addition of the ECL chemiluminescent reagent (Merck-Millipore). The light generated was measured using Spectramax M3 plate reader at a wavelength of 700 nm.

### Competitive enzymatic immunoassay

The presence of neutralizing antibodies was determined using the enzyme immunoassay adapted from NeutraLISA (EUROIMMUN, Brazil, kindly donated by Dr. Michael Soane). The kit contained a plate sensitized with SARS-CoV-2 S1 protein. One hundred 100 µL of a mixture of the test serum (final dilution 1/5) and the kit ACE2-biotin solution were added. After one hour at 37 °C, the wells were washed with the kit wash solution and incubated with the kit streptavidin-peroxidase solution for an additional 30 minutes at room temperature. The wells were washed again, and binding was detected by incubation with the chromogen contained in the kit for 15 minutes after the reaction was terminated by the addition of the kit stop solution. Absorbance values were then measured at 450 nm using the SpectraMax M3 plate reader. The same method was adapted by coating 100 µL of high binding transparent polystyrene plates with 50 ng of the recombinant RBD protein, as described in the ELISA assay. The next day, the wells were washed with the solution provided in the kit and the procedure was repeated as described above. To obtain the blank value, only the solution containing biotinylated ACE2 was used. Assay values were plotted as % inhibition using the following calculation: 100% -(sample value x 100/blank value) = % inhibition.

### Virus neutralization test

Cytopathic effect SARS-CoV-2 virus neutralizing tests were performed as described earlier [31-33] to quantify neutralizing antibody titers. For this, monolayers containing 5 × 10^4^ Vero cells (ATCC CCL-81) were grown in 96-well plates and exposed to Tissue Culture Infectious Dose (TCID_50_) of the Wuhan WT strain (SARS-CoV-2/human/BRA/SP02/2020 strain -M T126808.1), Delta variant (gisaid EPI_ISL_2965577), Gamma variant (gisaid EPI_ISL_1060981), and Omicron variant (hCoV-19/Brazil/SP-HIAE-ID990/2021 gisaid EPI_ISL_6901961) all previously incubated with 1:40 – 1:2560 serum. After 72 hours of incubation, all wells were evaluated by optical microscopy for the presence of cytopathic effects, a characteristic of SARS-CoV-2. Absence of cytopathic effects in at least the 1:20 dilution sample was considered a positive result of neutralizing antibodies for SARS-CoV-2. All cytopathic virus neutralization test (CPE-VNT) procedures were performed at a biosafety level 3 in the laboratory of the Federal University of São Paulo, according to World Health Organization recommendations.

### Statistical analysis

In vivo experiments were performed with a sample number of 5 animals per group and analyzed in duplicates from each animal. Results were expressed as mean +/-standard deviation. Statistical tests were analyzed using GraphPad Prism 9 software, with significant differences considered when p < 0.05 with ANOVA test with Tukey’s multiple comparison or Student’s t-test.

## Results

### Expression of the recombinant protein RBD in HEK293 cells

We have used the HEK293 cell line, whose cells are widely employed to produce recombinant proteins because they have the machinery capable of performing much of the folding and post-translational processing to produce functional proteins derived from mammalian and non-mammalian DNA sequences [34]. The cells were transfected with pcDNA 3.1/RBD plasmid, and after selecting cells resistant to geneticin G418, the cells were cloned. We first used an immunofluorescence assay to detect reactivity with anti-RBD antibodies before cloning (**Fig**. 1A). These were later used for detecting positive clones. One of these clones was used to check the expression of RBD protein by Western blotting. As expected, HEK293 cells expressed and secreted the RBD in the supernatant (**Fig**. 1B). The expected molecular mass for the recombinant protein is 28 kDa and a band with 37 kDa suggests that it was glycosylated.

**Fig. 1.**
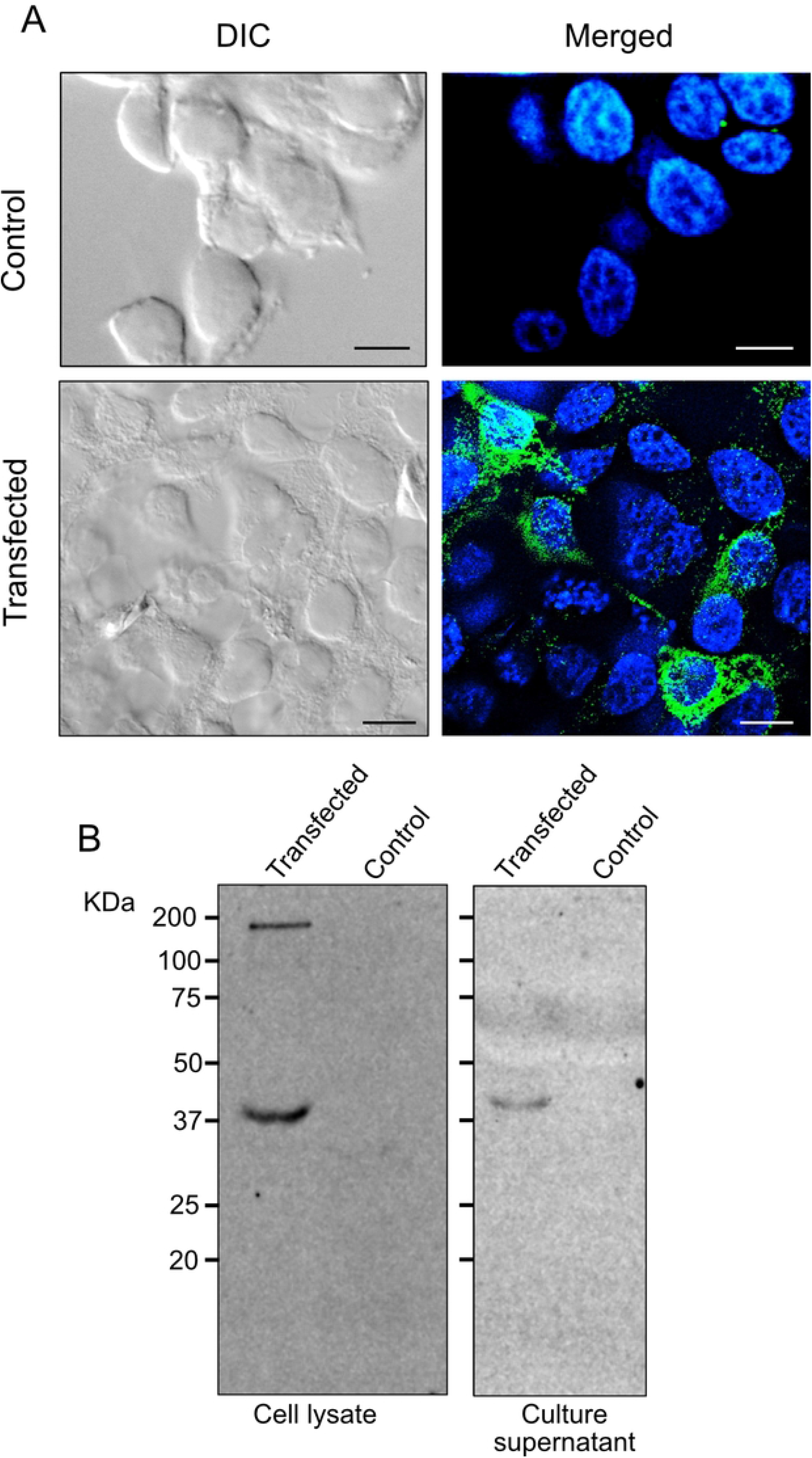
RBD is expressed and secreted by HEK293 cells. **A**. Immunofluorescence of control and transfected HEK293 cells after G418-selection cells using anti-mouse RBD antibodies (green) and DAPI staining (blue). The images show DIC and merged fluorescent images. Bars = 5 µm. **B**. Western blotting of cell lysates and culture supernatant collected after 24 hours of culture of HEK293 cells containing the RBD plasmid and non-transfected cells revealed with horse serum anti-RBD and peroxidase-conjugated protein A. The size markers in kDaltons are shown on the left.

The RBD protein secreted in the supernatant of HEK293 cells was initially purified on a Ni-Sepharose chromatography column, as it contained a histidine tag in the C-terminus. **Fig**. 2A shows an example of the results obtained after Ni-Sepharose purification. The fractions eluted from the Ni-Sepharose column with 0.5 M imidazole were concentrated and subjected to a new purification step on a Superdex 200 column. Some protein eluted in the column void with a mass greater than 250 kDa, and two additional peaks were observed (**Fig**. 2B). One corresponded to a mass of 135 kDa and another of about 160 kDa. All these samples contained the recombinant protein RBD that migrated in the SDS-PAGE with 37 kDa (inset). The band eluted in the void also contained a lower amount of the 37 kDa protein suggesting that it may correspond to protein aggregates due to its higher light absorption at 280 nm. The fractions eluting at 135 kDa were pooled, concentrated, and re-applied in the Superdex-200 column. Indeed, they eluted mainly in the column with an elution volume that matched the size of 125 kDa (**Fig**. 2C). In summary, we obtained 2.8 mg of protein in trimeric form from 1 L of supernatant of HEk293 cell culture.

**Fig. 2.**
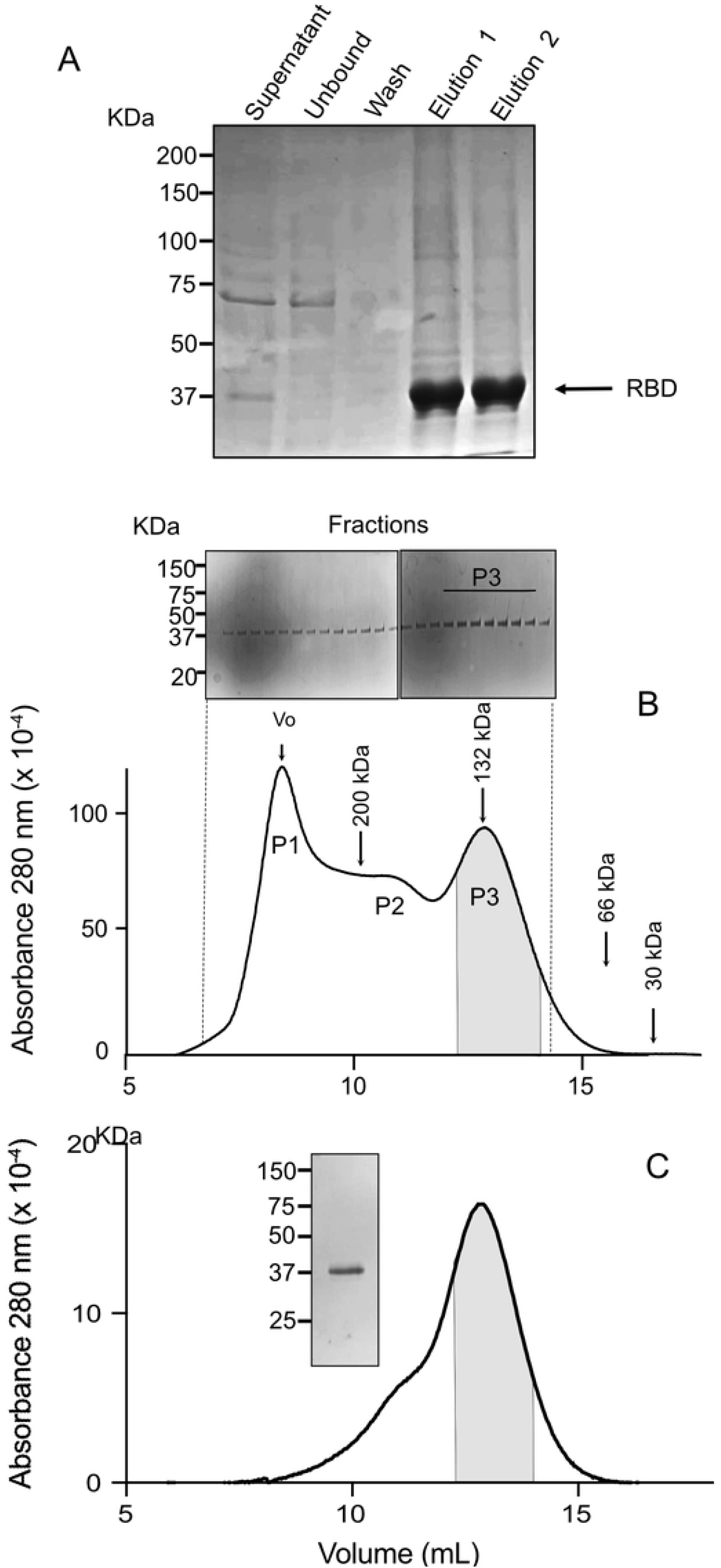
RBD purification steps. **A**. SDS-PAGE Gel stained with Coomassie blue R250 of a 20 µL of samples of a total of 400 mL of culture supernatant applied in the Ni-Sepharose column (Supernatant). The fractions corresponding to unbound and washed eluates from the column and the two eluates (5 mL each) in buffers containing 0.5 M imidazole are indicated. **B**. Size exclusion chromatography profile of the Superdex 200 column. The fractions concentrated material eluted from the Ni-Sepharose column were concentrated to 0.5 mL and applied in the column. Proteins were detected by absorbance at 280 nm. P1 corresponds to possible protein aggregates, P2 to protein forms with a mass higher than 200 kDa, and P3 to a mass of 125 kDa. The inset shows Coomassie bluer R250 stained SDS-PAGE of the corresponding fractions eluted from the column. The arrows indicate the elution position of the size markers (β-amylase, 200 kDa; BSA dimer, 132 kDa; BSA monomer, 66 kDa and carbonic anhydrase, 30 kDa. **C**. Elution profile of concentrated samples of the first exclusion chromatography (gray area in **B**) in a new chromatography. The inset shows a typical SDS-PAGE gel stained with Coomassie Blue R250 of a concentrated pool of fractions as shown in the gray area. Size markers for SDS-PAGE are indicated in kDa.

### The purified RBD formed multimeric structures

Because the protein eluted with approximatively 135 kDa from the size column, we performed an SDS-PAGE in the absence of the reducing agent. We found that 2/3 of the protein migrates with a mass of 75 kDa and 1/3 with a mass of 37 kDa (**Fig**. 3A), suggesting that the trimeric form was formed by a dimer of two covalently linked monomers by at least one disulfide bond and one monomer. This may be explained by the presence of 9 cysteines in each monomer of our construction. The presence of a higher mass band could correspond to structures of large size consistent with the second peak in the elution profile of a band above 150 kDa in the Superdex-200 column. To verify whether these structures are formed, we treated the purified protein with BS3, a crosslinking reagent that couples close free amino groups. We found two major bands with 130 kDa and approximatively 260 kDa (**Fig**. 3B), corresponding to three and six units of the original protein. We also confirmed that the RBD multimers contained glycosidic residues by reactivity with concanavalin A (**Fig**. 3C) and displayed a folded structure as seen by circular dichroism analysis (**Fig**. 3D), with similar levels of α and β sheets as seen in monomeric RBD [35].

**Fig. 3.**
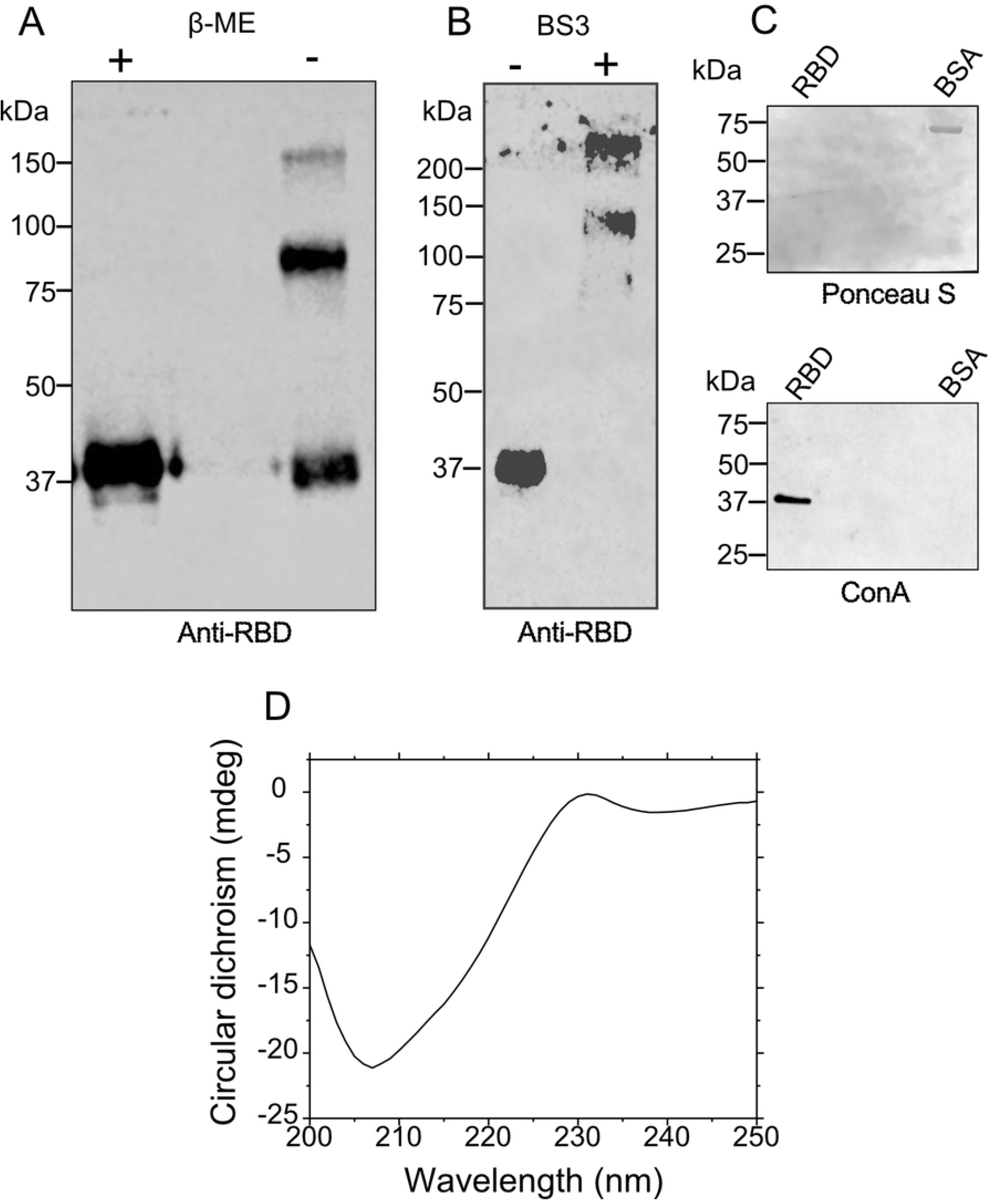
The recombinant RBD forms a trimeric structure and presented a structured conformation. **A**. The purified protein eluted from the second sizing column was concentrated and boiled in sample buffer in presence (+) or absence (-) of β-mercaptoethanol (βME) and submitted to Western Blotting using mouse anti-RBD antibodies. **B**. The purified RBD sample was treated in the absence (‾) or presence (+) of the BS3 crosslinker for 1 hour and submitted to Western Blotting using mouse anti-RBD. **C**. A nitrocellulose membrane obtained after the transfer of a SDS-PAGE loaded with the recombinant RBD or BSA and stained with Ponceau S (Top), or incubated with biotin-Concanavalin A (ConA) and revealed with streptavidin-peroxidase by ECL reaction. **D**. Circular dichroism analysis of the purified RB at 0.3 mg/mL in TBS.

### The multimeric RBD induced high titers of antibodies in mice

To evaluate the antigenicity of the multimeric RBD, mice were immunized intramuscularly with three doses containing 100 µg of the recombinant RBD mixed with aluminum hydroxide at two-week intervals. Control animals received only PBS with the adjuvant. At each administration, 10 µl of blood was collected from each animal to analyze antibody production. The first dose of the immunogen did not stimulate the production of detectable level of antibodies when the serum was diluted 1/100 in a chemiluminescent ELISA assay with plates containing recombinant RBD (**Fig**. 4A). A response was detected after the second dose and further increased after a third injection (**Fig**. 4A). We then checked the titers of the pooled antisera obtained in ELISA assays. In this experiment, we used 12.5 to 400 ng of the multimeric RBD to coat each well of the plates and we observed binding of the sera at dilutions up to 1:50.000 (**Fig**. 4B). No reactivity was found in the pre-immune sera at this dilution.

**Fig. 4.**
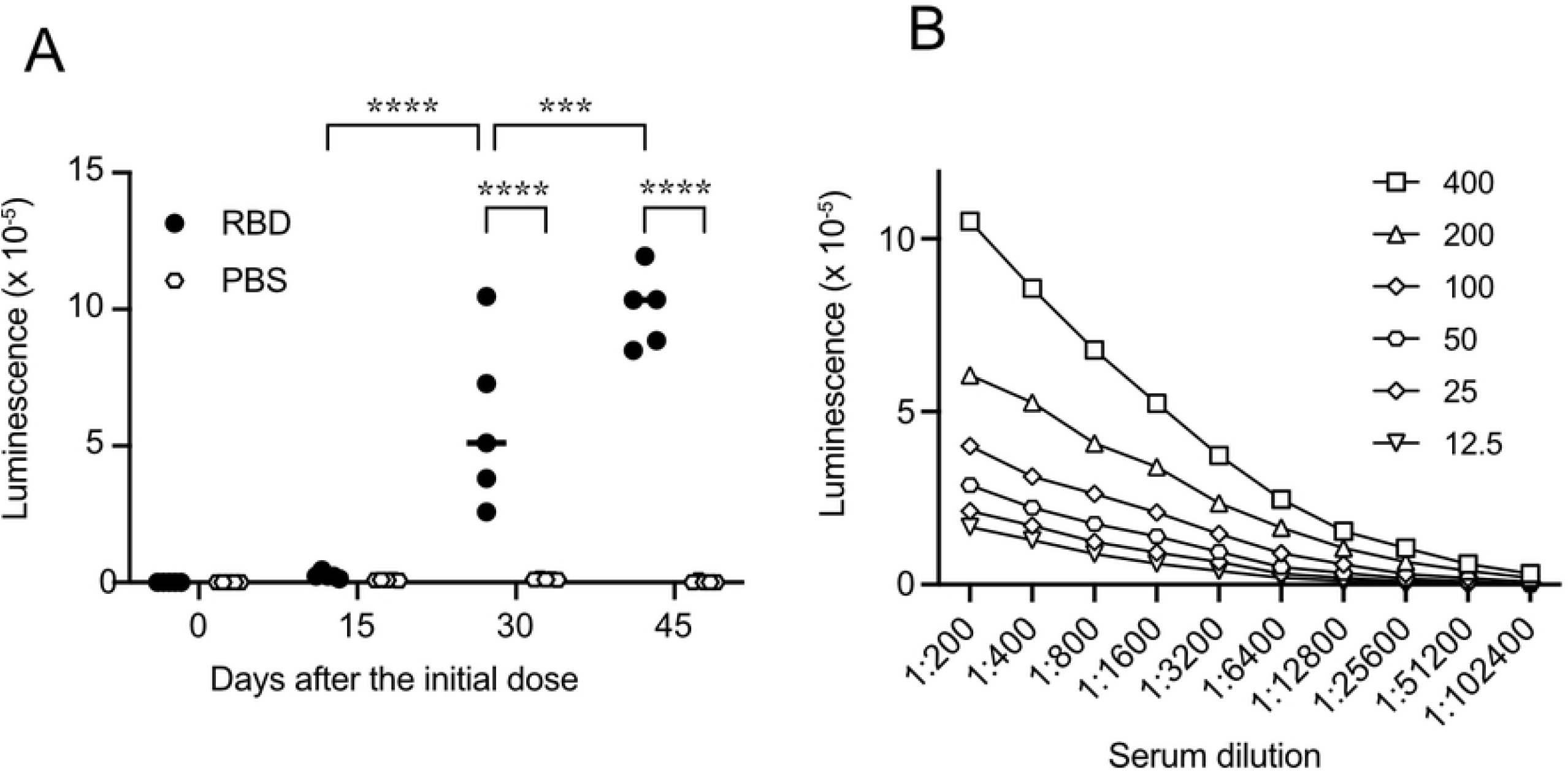
ELISA assay used to evaluate the reactivity of sera to the multimeric RBD. **A**. The purified multimeric RBD (200 ng), was used to coat an ELISA plate. Assays were performed in duplicate with sera diluted 1:100. Bound antibodies were detected with peroxidase-coupled anti-mouse IgG antibodies and visualized with ECL. Signals correspond to the mean of duplicates of sera obtained from RBD immunized mice (closed circles) or PBS (open lozenges). The significance of the indicated differences was calculated using the two-way test ANOVA with Tukey’s multiple comparison test, where **** represents p < 0.0001 and *** represents p < 0.001. **B**. Determination of the titer of pooled sera obtained after the third immunization in plates coated with the indicated amounts of RBD per well. Points represent mean of duplicates for each dilution of sera incubated in PBS with 3% BSA.

### Analysis of antibody neutralization by competition assay with ACE2

To analyze whether the resulting sera can neutralize the binding of RBD to ACE2, we performed a competitive ELISA with the immobilized S1 glycoprotein followed by addition of ACE2-biotin. In this assay, we found that all five mouse sera could inhibit the binding of ACE2 to the viral protein comparable to a positive control containing a neutralizing antibody (**Fig**. 5A). We also observed that ACE2-biotin binds to wells coated with different concentrations of the multimeric RBD adsorbed to the plate (**Fig**. 5B). The interaction follows a saturation curve. Therefore, the wells were coated with 50 ng of RBD, which was the concentration in the linear range of the binding assay, and then used to check for antibody inhibition diluted 1:5. Indeed, all sera prevented the binding of ACE2 with three of them being more effective (**Fig**. 5C). When tested for virus neutralization. Two mouse sera immunized with multimeric RBD protected against the SARS-Cov-2 infection (Wuhan WT strain) at 1:80 dilution and one at 1:20 (**Fig**. 5D). These sera coincided with those shown with higher inhibition of ACE2 binding. The two remaining sera were not able to protect infection at least at 1:20 dilution. The protective antibodies could not prevent infection by three other SARS-CoV-2 strains at least up to the dilutions used.

**Fig. 5.**
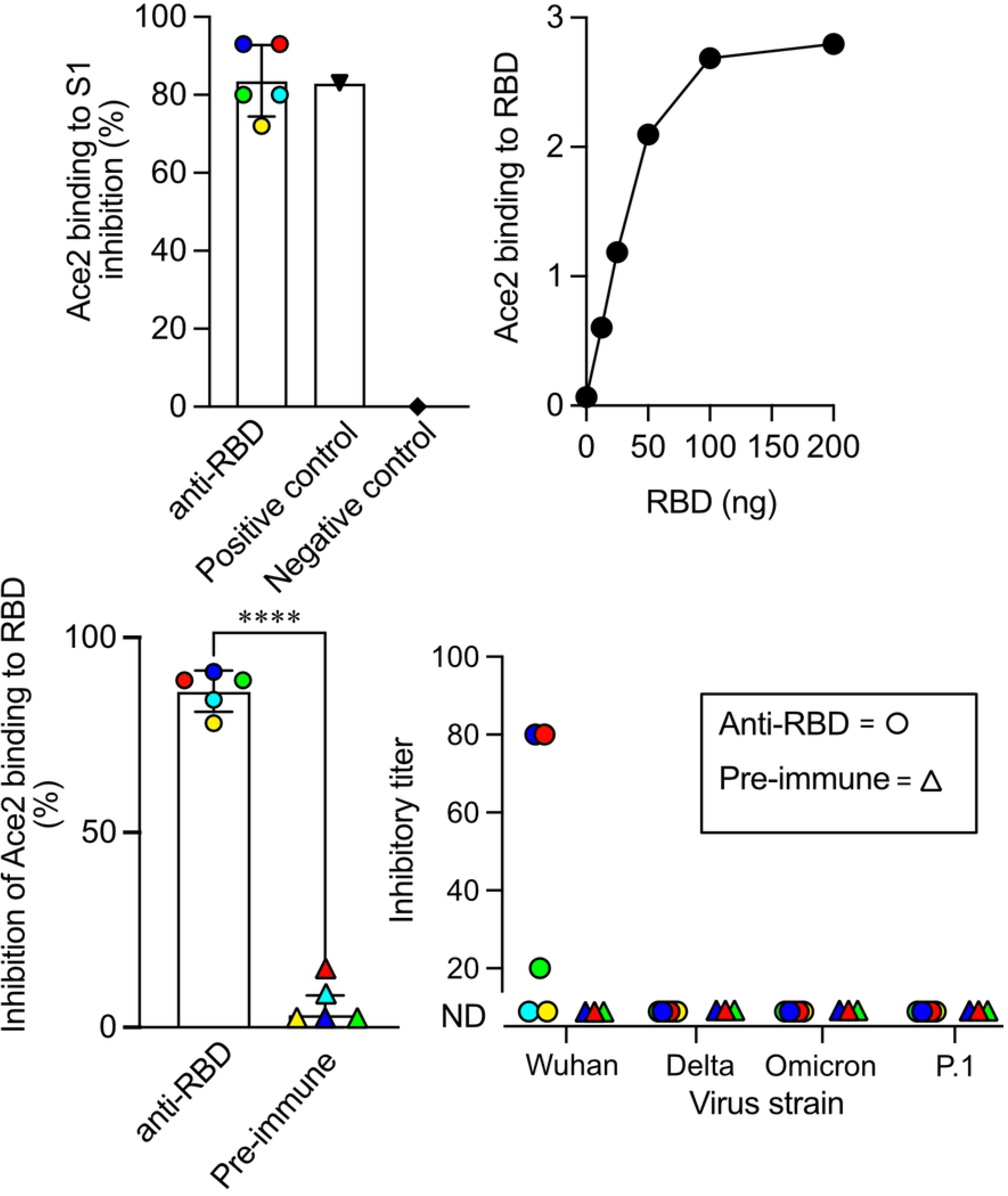
Mouse antibodies block ACE2 binding to the multimeric RBD and prevent virus infection. **A**. Competition assay of ACE2 binding to immobilized S1 protein. Wells were incubated with the ACE2-biotin solution in the presence of the positive (+) and negative (-) control provided with the kit and with mouse sera collected after the third immunization. **B**. ACE2-biotin protein binding to wells coated with the indicated amount recombinant RBD in 50 µL of 0.1 M sodium bicarbonate buffer, pH 8. After washing, the wells were incubated with streptavidin solution coupled to peroxidase and associated peroxidase measured by the Immunogen detection system. **C**. In vitro neutralization assay of ACE2-biotin binding to wells coated with 50 ng of multimeric RBD in the presence of 1:5 diluted mouse anti-RBD or the respective pre-immune serum. Student’s t-test was used to compare the groups, where **** represents p < 0.0001. **D**. Titers of each mouse serum for the inhibition of SARS-CoV-2 infection of Vero Cells. The immune sera are indicated by circles and pre-immune by triangles. The colors in A, C and D show the corresponding sera.

## Discussion

The search for information on the structure and mechanism of the SARS-CoV-2 virus has undoubtedly reached an extraordinary pace. Researchers around the world have spared no effort to gain knowledge and this has led to very effective diagnostic methods and vaccines in a very short time. Today, thanks to the effectiveness of the vaccines already produced, the pandemic is under control, although treatment options for the disease remain limited and we do not know the level of protection existing vaccines will provide against new viral mutations. The new variants that have emerged are due to mutations in the RBD region of S1. Although immunization with more than three doses of the already available vaccines has been shown to elicit neutralizing antibodies that are approximately 80% protective against the new variants, it is unknown to what extent this will be maintained [36-39].

In this work, we generated a new recombinant RBD with the full carboxy terminus of the S1 virus subunit and analyzed its ability as a protective antigen upon immunization. This new recombinant protein was produced and released constitutively by HEK293 cells that behaved like multimeric structures, most likely trimers and hexamers. Some of the protein that was produced formed multimeric aggregates of high molecular mass, which could be removed by gel exclusion chromatography. Our analyses revealed that the trimer consists of a covalent dimer connected by an S-S bridge, which is consistent with the fact that there is a cysteine in the C-terminal region in our construct. In the protein structure, this cysteine is far from the ACE2 binding site, while other cysteines form disulfide bridges within the chain, which are important for stabilizing the RBD structure [40]. We do not yet know whether the trimeric or hexameric structures have a similar conformation to the trimer formed by the S1 protein on the surface of the virus. The circular dichroism showed homogenous folding with 52.5% alpha-helix, 18.5% beta sheets and 5.5% coils in a similar manner to published spectra of monomeric RBD and reinforces the notion that in our case each monomer acquires a conformation similar to that found in the virus.

We verified that the protein immobilized on a polystyrene plate was recognized by the biotin labeled ACE2 receptor. This recognition takes place through the RBM region located between waste 438 to 508 [41]. This region is exposed in the Virus Protein S1 after cleavage with furin if raised on the trimer to recognize the receptor [42]. In addition, in the trimer, only one RBD binds to ACE2 in the membrane at any one time, due to the impossibility of coupling each receptor molecule to the three subunits by conformational impediment [43].

The amino acid sequence of the recombinant protein (**Fig**. S2) has two N-glycosylation regions, at asparagine 12 and 24. The sequence also has an RGD-motif that binds to integrins that may be co-receptors for SARS-CoV-2 to infect cells [44]. The expressed protein was also recognized by the lectin Concanavalin A, a mannose-binding lectin [45], in agreement that mannose is one of the main carbohydrates present in the structure of the Spike protein [46]. In fact, some studies have shown the importance of glycosylation for the interaction of RBD-ACE2, in addition to the involvement in protein stability and functionality [47]. Furthermore, the interaction of the Spike glycoprotein with receptors (DC/L-SIGN) that bind to carbohydrates that are present in the cell membrane, suggested that the glycosylated part of the protein is important in viral invasion into cells [48].

The cells generated in our study are stable and can be adapted to grow on a large scale. We also developed an ELISA/ECL assay using the recombinant multimeric RBD protein, which can be very useful for identifying antibodies in the human population. This is because we can continuously obtain a large amount of recombinant protein with our stable system without having to repeat transfection for each new purification, and the assay can be adapted to characterize the presence of different classes and types of antibodies. To evaluate the potential of RBD as an antigen, we immunized BALB/c mice with three doses. The sera obtained after the third dose immunization of mice with Alum as adjuvant showed very high antibody titers and contained antibodies that were able to inhibit binding to ACE2 to the full S1 chain or the multimeric RBD. Furthermore, three out of five mouse sera, were able to neutralize the infection of Vero Cells by SARS-CoV-2 at relevant dilutions (above 1:20). These findings open perspectives to produce protein to be tested in other animal models and eventually in the human population, either as a single or combined vaccine.

Neutralizing antibodies are known to target the receptor-binding domain, which is 90% neutralizing in convalescent sera [49], supporting the use of RBD as an antigen for vaccines. Studies published early in the pandemic demonstrated that a DNA vaccine containing the RBD along with a trimerization tag was able to elicit a protective immune response in a model of *rhesus macaques* [50]. It has been shown that trimeric RBD interacts with greater affinity with ACE2 receptor molecules on the cell surface than monomeric RBD [51]. Therefore, production of antibodies that recognize the trimeric structure may be more effective in inhibiting this binding. This is related to the fact that immunogens with a more multimeric structure elicit a more robust humoral response in mice [52]. In addition, proteins based on total protein S may exhibit some difficulties in stabilizing the prefusion conformation [53] and may fail to generate adequate antibodies.

The antibodies developed in mice with the multimeric RBD protein failed to prevent invasion at dilutions above 1:20 of anything other than the SARS-CoV-2 Wuhan WT strain. Although several studies have shown cross-reactivity among HCoVs, as they contain conserved regions [54], especially in vaccines, mAbs isolated before the appearance of variants showed variable protective capacity [55]. For example, antibodies generated by the Alpha, Beta, Gamma, and Delta variants are not efficient against Omicron, even though they share some mutations [56]. In contrast, it has been shown that monoclonal antibodies made from a recombinant RBD (Wuhan sequence) neutralized major variants [57]. However, in our case a full C-terminal domain of the S1 protein along the RBD domain was introduced and might affect the production of new neutralizing antibodies. In fact, some publications indicate that the full extension of S1 participates in the interaction of the virus with the ACE2 receptor [58] and the generation of a robust inhibition of ACE2 binding may suggest that this portion also exerted a role in the generation of neutralizing antibodies in our case. This, together with the decrease in antibody titers after immunization, clearly demonstrates the requirement of timely vaccine boosts with new variants [59]. In addition, when SARS-CoV-2 binds to ACE2, it reduces its expression in infected cells [60]. It would therefore be interesting to test whether the antibodies produced would also be able to alter internalization via alternative receptors.

Although there are already several constructs using trimeric RBD as an antigen, most of them do not use the extended (residues 331 to 589) portion of the S1 protein C-terminal domain [17, 18, 51, 61, 62]. Usually, the RBD domain expands the protein to the residue 527. The presence of the expansion to residue 541 in SARS-CoV-2-CTD enhances the binding to ACE2 and affects the immunogenicity of the protein [63]. The extension we introduced (549-589) is exposed outside the trimeric structure and might help to fold the RBD-CTD and allow further recognition by antibodies.

conclusion, this study describes the production of a new RBD multimeric protein that can act as an efficient antigen for diagnosis and neutralization tests. In addition, it is an alternative subunit vaccine for SARS-CoV-2 that can be used as a platform for construction of immunization variants. Our observations also support previous studies that have shown the efficacy of multimeric RBD. Although there are already several constructions using RBD as an antigen, in this version we described an extended C-terminal and a foldon domain generating a new multimeric form. We are still living in the end stage of a pandemic. SARS-CoV-2’s ability to mutate is well known and there is a lack of antivirals, so this is a welcome addition to the range of tools that combat the disease.

## Author contributions

Conceptualization: V.A.de L, P.L.H, S.S; Methodology: V.A. de L., R. da S.F., J.T.M; Formal analysis: V.A. de L., S.S.; Investigation: V.A.de L., R.A.S., J.T.M.; Data curation: S.S.; Writing - original draft: V.A.de L., S.S.; Writing - review & editing: S.S., M.L.V.O, J.T.M, P.L.H.; Visualization: V.A. de L., M.Z., N.L.C.S., C.L.A.; Supervision: S.S.; Project administration: S.S.; Funding acquisition: S.S.,M.L.V.O, P.L.H., L.M.R.J.

## Data availability

All data and resources described in this manuscript are available upon request.

## Abbreviations

RBD: receptor binding domain
ACE2: angiotensin 2 converting enzyme
CPE-VNT: cytopathic virus neutralization test.

## Supplementary material

**Fig. S1.**
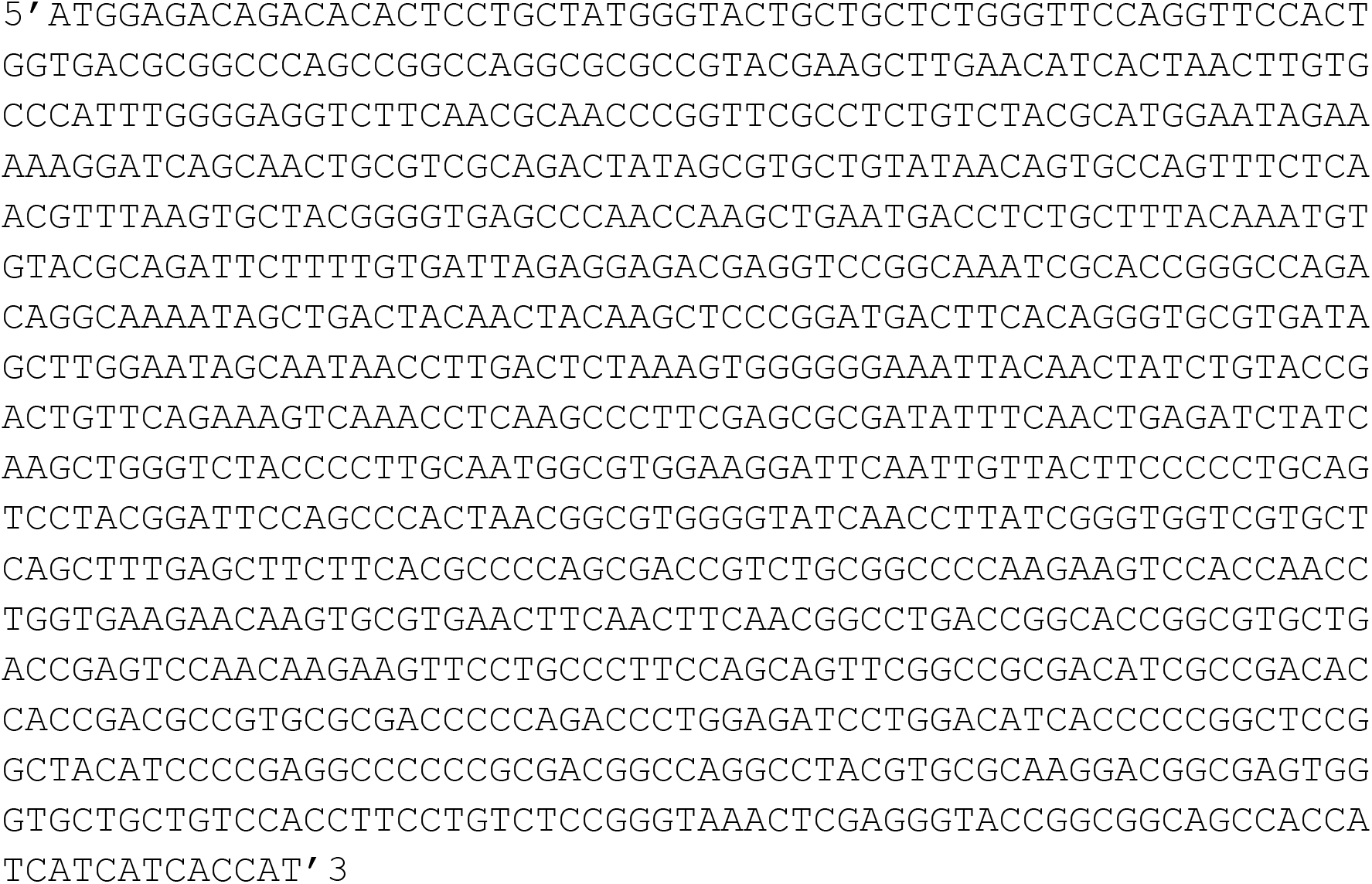
Sequence inserted into the pcDNA4 plasmid.

**Fig. S2.**
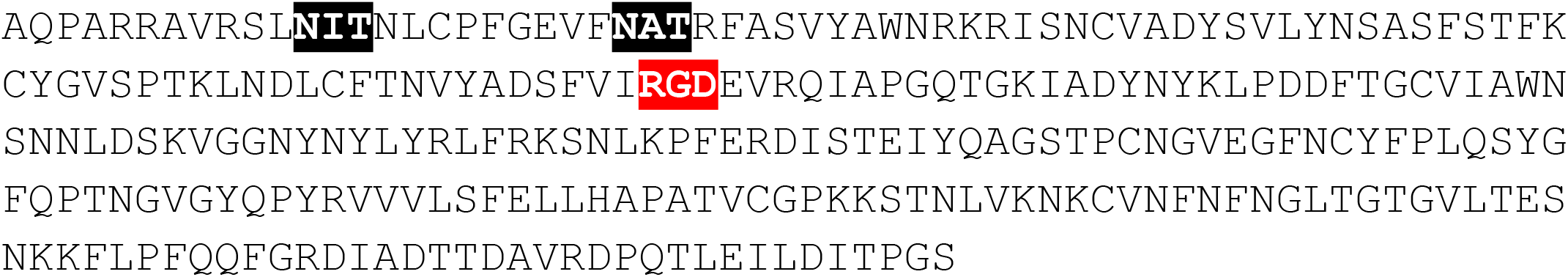
Amino acid sequence of the recombinant protein. Highlighted in black N-glycosylation region. In red, integrin-binding motif sequence.

